# A computational pipeline to determine lobular electric field distribution during cerebellar transcranial direct current stimulation

**DOI:** 10.1101/531699

**Authors:** Zeynab Rezaee, Anirban Dutta

## Abstract

**Objective:** Cerebellar transcranial direct current stimulation (ctDCS) is challenging due to the complexity of the cerebellar structure. Therefore, our objective is to develop a freely available computational pipeline to perform cerebellar atlas-based electric field analysis using magnetic resonance imaging (MRI) guided subject-specific head modeling.

**Methods:** We present a freely available computational pipeline to determine subject-specific lobular electric field distribution during ctDCS. The computational pipeline can isolate subject-specific cerebellar lobules based on a spatially unbiased atlas (SUIT) for the cerebellum, and then calculates the lobular electric field distribution during ctDCS. The computational pipeline was tested in a case study using a subject-specific head model as well as using a Colin 27 Average Brain. The 5cmx5cm anode was placed 3 cm lateral to inion, and the same sized cathode was placed on the contralateral supraorbital area (called Manto montage) and buccinators muscle (called Celnik montage). A 4×1 HD-ctDCS electrode montage was also implemented for a comparison using analysis of variance (ANOVA).

**Results:** Eta-squared effect size after three-way ANOVA for electric field strength was 0.05 for lobule, 0.00 for montage, 0.04 for head model, 0.01 for lobule*montage interaction, 0.01 for lobule* head model interaction, and 0.00 for montage*head model interaction in case of Enorm. Here, the electric field strength of both the Celnik and the Manto montages affected the lobules Crus II, VIIb, VIII, IX of the targeted cerebellar hemispheres while Manto montage had more bilateral effect. The HD-ctDCS montage primarily affected the lobules Crus I, Crus II, VIIb of the targeted cerebellar hemisphere. Our freely available computational modeling approach to analyze subject-specific lobular electric field distribution during ctDCS provided an insight into healthy human anodal ctDCS results

## 1. Introduction

Motor adaptation is critical for performing activities of daily living where we need to continuously adjust our movement to account for the changes in the environment. This is based on sensory feedback and motor skill learning to perfect our intended movement pattern in an ever-changing environment. One kind of sensorimotor learning involves building a specific internal model of the environment for feedforward prediction during motor control (Thoroughman and Shadmehr, 1999) to overcome time delays associated with feedback control (Wolpert et al., 1998). An important application of such learning is to compensate for environment perturbations where Huang and colleagues (Huang et al., 2011) suggested that the motor memory has three components, a feedforward prediction model, a bias in motor output due to task repetition, and a reinforcement-dependent bias due to reward dependent association (Vaswani and Shadmehr, 2013). Here, cerebellar architecture has been found to support the computations required by the feedforward prediction model from animal studies as well as from studies on patients with cerebellar dysfunction (Ebner, 2013). Marr-Albus-Ito hypothesis is the earliest and most studied mechanism that is based on the architecture of the cerebellar cortex and assigns specific functions to the climbing fiber-Purkinje cell and the mossy fiber-granule cell-parallel fiber-Purkinje cell circuits (Popa et al., 2016). A significant concentration of neurons in cerebellum is due to densely packed small granule cells (van Dun et al., 2016) that can support matrix memory based on Hebbian learning primarily for 'readout tables'. Cerebellum has an essential role in motor learning and coordination as well as non-motor functions such as sensory and cognitive processes (Buckner, 2013). Here, neuroplasticity can be facilitated with an extrinsic stimuli using noninvasive brain stimulation (NIBS) (Hallett, 2005), which has been shown recently to modulate brain (Shafi et al., 2012) and spinal (Wolpaw, 2010) network interactions. Specifically, Purkinje cell firing has several of the characteristics of a forward internal model (Ebner, 2013) which is the main target of cerebellar transcranial direct current stimulation (ctDCS) (Galea et al., 2009) - a painless non-invasive technique where a weak direct current (i.e., up to 2mA) is delivered through a scalp electrode overlying the cerebellum (van Dun et al., 2016).

The ctDCS technique has been shown to be safe in humans with no adverse side effects; however, its efficacy appears to be limited (Ferrucci et al., 2016). Here, the modulation of the activity in the cerebellar neurons (cerebellar excitability) with the electric field is the goal (Ferrucci et al., 2015), but it is very challenging due to the extreme folding of the cerebellar cortex. Galea and colleagues proposed that ctDCS produces polarity specific effects by polarizing the Purkinje cells, thereby affecting the activity in the deep cerebellar output nuclei (Galea et al., 2009). Cerebellar role in modulating sensory processing has also been demonstrated (Popa et al., 2013), which can explain the ctDCS effects on distant plasticity in human cortical areas (i.e., the motor cortex) (Grimaldi et al., 2016). Indeed, cerebellum serves two purposes, i.e., Purkinje cells provide both the internal prediction and sensory feedback signals for motor control (Popa et al., 2016). Furthermore, optimization of the lobular electric field distribution to target cerebellar motor syndrome and/or cognitive performance is important (Stoodley and Schmahmann, 2009). Therefore, a systematic investigation of subject-specific lobular electric field distribution based on a cerebellar atlas is necessary to delineate the direct electric field effects on the cerebellum from extra-cerebellar effects. Here, computational modeling studies have shown that the ctDCS electric field distribution reaches the human cerebellum where the posterior and the inferior parts of the cerebellum (i.e., lobules VI-VIII) are particularly susceptible (Grimaldi et al., 2016). However, a lobular atlas-based analysis of subject-specific electric field distribution during ctDCS was not found in the literature (Priori et al., 2014),(Parazzini et al., 2014),(Fiocchi et al., 2016). Therefore, the main objective of this technology report is to present a freely available computational pipeline that allows visualization of the lobular electric field distribution during ctDCS. This will facilitate optimization of the lobular electric field distribution which is important to specifically target cerebellar motor syndrome (anterior lobe, extending into VI) versus cognitive performance (posterior lobe, including lobules VII and VIII) (Stoodley and Schmahmann, 2009).

The challenges in lobular targeting of ctDCS includes high conductivity of the cerebrospinal fluid (CSF) and extreme folding of the cerebellar cortex (Fiocchi et al., 2017). Here, the goal is an optimal electrode placement, e.g., with more focal high-definition (HD) ctDCS montages (Fiocchi et al., 2017), that can in principle direct the electric field towards deeper targets by taking advantage of the high conductivity of the CSF and the interhemispheric fissure. Moreover, computational modeling (Priori et al., 2014),(Parazzini et al., 2014),(Fiocchi et al., 2016) is important to address the inter-subject variability in the ctDCS effects for its clinical translation (Ferrucci et al., 2016). This is crucial since ctDCS effects were recently said to be mediated by mechanisms other than cerebellar excitability changes (Grimaldi et al., 2016) since non-focal electric field with two-electrode montages may affect brain areas other than cerebellum. In this technology report, we aimed to develop the computational modeling pipeline using freely available software packages that are easily accessible worldwide to facilitate clinical translation of ctDCS. Therefore, we developed a modeling pipeline that creates a head model from subject’s magnetic resonance imaging (MRI) and simulates the ctDCS field distribution in cerebellar lobules using a freely available software (Opitz et al., 2015) - SimNIBS. Our main contribution is in providing an approach for the isolation of the cerebellum and its lobules based on Spatially Unbiased Infratentorial Template for the Cerebellum (SUIT) atlas (Diedrichsen et al., 2009) which allowed us to investigate the lobular electric fields following finite element analysis (FEA) in SimNIBS. We have adapted the SUIT isolation and activation visualization scripts, which are commonly used to analyze functional MRI activation maps, to analyze the lobular electric fields due to ctDCS. Our SUIT–based approach to determine cerebellar lobular electric field distribution can be applied to FEA results from different freely available as well as commercially available software (e.g., COMSOL). To show that visualization of cerebellar lobular electric field distribution can provide further insights, we applied our pipeline to analyze our healthy human experimental results during visuomotor learning of myoelectric visual pursuit using electromyogram (published in a conference (Abadi and Dutta, 2017)).

## 2. Material and Methods

We developed the computational modeling pipeline using freely available software packages that are easily accessible worldwide to facilitate clinical translation of ctDCS. Using our computational modeling pipeline, we investigated two common ctDCS montages (Grimaldi et al., 2016),(Grimaldi et al., 2014) with the anode placed over the right cerebellum, and **1.** the cathode placed over the right buccinator muscle – called Celnik montage henceforth, **2.** the cathode placed on the contralateral supraorbital area – called Manto montage henceforth. We also investigated a recently published high-definition (HD) ctDCS 4×1 montage (Doppelmayr et al., 2016). Our computational pipeline leverages SUIT, which is one of the automated algorithms developed explicitly for cerebellum segmentation (Diedrichsen, 2006), and is a freely available SPM (Statistical Parametric Mapping (SPM - Statistical Parametric Mapping)) toolbox for functional MRI data analysis. In this toolbox, a probabilistic atlas of the cerebellar nuclei, a cerebellar cortical parcellation atlas in MNI (Montreal Neurological Institute) space, and SUIT template are available. Therefore, we used SUIT SPM toolbox for isolation of cerebellar lobules where SUIT provided an improved and fine-grained exploration, registration and anatomical detail of the cerebellum for structural and electric field images. Since our SUIT–based approach to determine cerebellar lobular electric field distribution can be applied to FEA results from different freely available FEA software so we compared the lobular electric field results based on FEA using SimNIBS pipeline (Opitz et al., 2015) with that using the Realistic vOlumetric-Approach to Simulate Transcranial Electric Stimulation (ROAST) pipeline (Huang et al., 2017).

### 2.1. Computational Modeling Pipeline

#### 2.1.1. MRI data acquisition and subject-specific head model creation

The first step in creating an anatomically accurate subject-specific head model is the segmentation of structural magnetic resonance images (MRI). The individual head model was constructed using MR images taken from a healthy volunteer in accordance with the Declaration of Helsinki - a statement of ethical principles for medical research involving humans. For research participation as well as for the publication of the case report including participant’s identifiable information, a written informed consent was obtained from the subject at the University at Buffalo prior to presenting this case. The subject did not have any history of neurological or psychiatric diseases. Images were taken from 3 Tesla Magnetic Resonance Imaging (MRI) system (Toshiba Vantage) at the University at Buffalo Clinical and Translational Science Institute using a sixteen multichannel receiver head coil. Two T1-weighted images (with and without fat suppression) were acquired for the subject (Windhoff et al., 2013). MR sequence consisted of the following parameters: MPRAGE, 192 slices, matrix size= 256 256, Flip/Flop angle=8/0, TR/TE=6.2/3.2. Also, two T2-weighted images (with and without fat suppression) were acquired for the subject with the sequence of 30 slices, matrix size of 256 256, flip/flop angle of 110/150 degree, and TR/TE=11990/108. From these four MR images, a tetrahedral volume mesh of the head was created using “mri2mesh” script which is provided in the SimNIBS package (Windhoff et al., 2013). The “mri2mesh” is based on four open source software; FreeSurfer (https://surfer.nmr.mgh.harvard.edu/), FSL (https://fsl.fmrib.ox.ac.uk/fsl/fslwiki), Meshfix (https://github.com/MarcoAttene/MeshFix-V2.1), and Gmsh (http://gmsh.info/). This script integrates all these software into a single pipeline for mesh generation from MR images (Windhoff et al., 2013). After segmentation using FSL and FreeSurfer, five tissues were modeled by the volume mesh; Skin, Skull, Cerebrospinal Fluid, Gray Matter, and White Matter. Different brain tissues for the volume mesh components were modeled as different volume conductors in SimNIBS with their own specific conductivity (Windhoff et al., 2013), as shown in Table 1. We also used Colin27 average brain (Holmes et al., 1998), which is the stereotaxic average of 27 T1-weighted MRI scans of the same individual, to create another head model for comparison (Colin27 head model has been published by Guhathakurta and Dutta – (Guhathakurta and Dutta, 2016)). The main contribution of this technology report is the freely available computational modeling pipeline to visualize subject-specific lobular electric field distribution following FEA, as shown in Figure 1.

**Table1.**
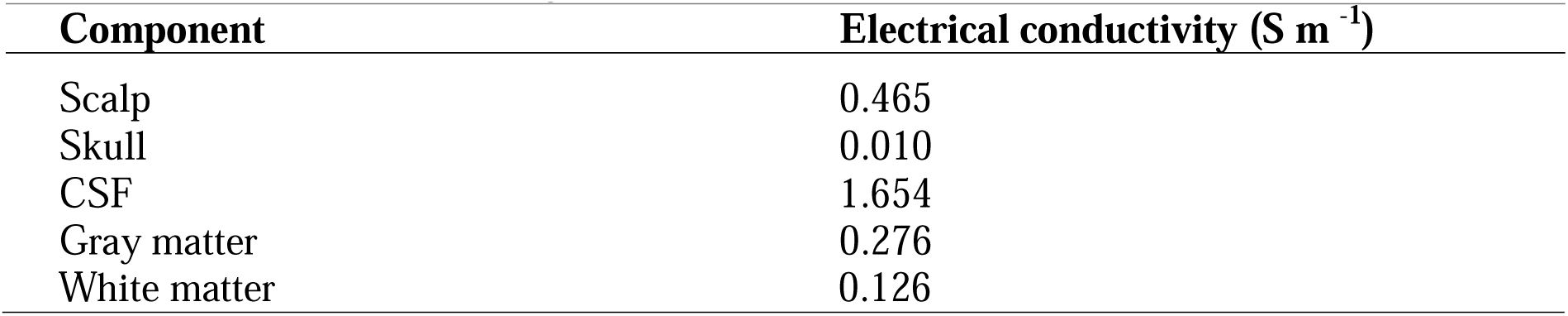
Electrical Conductivity

**Figure 1.**
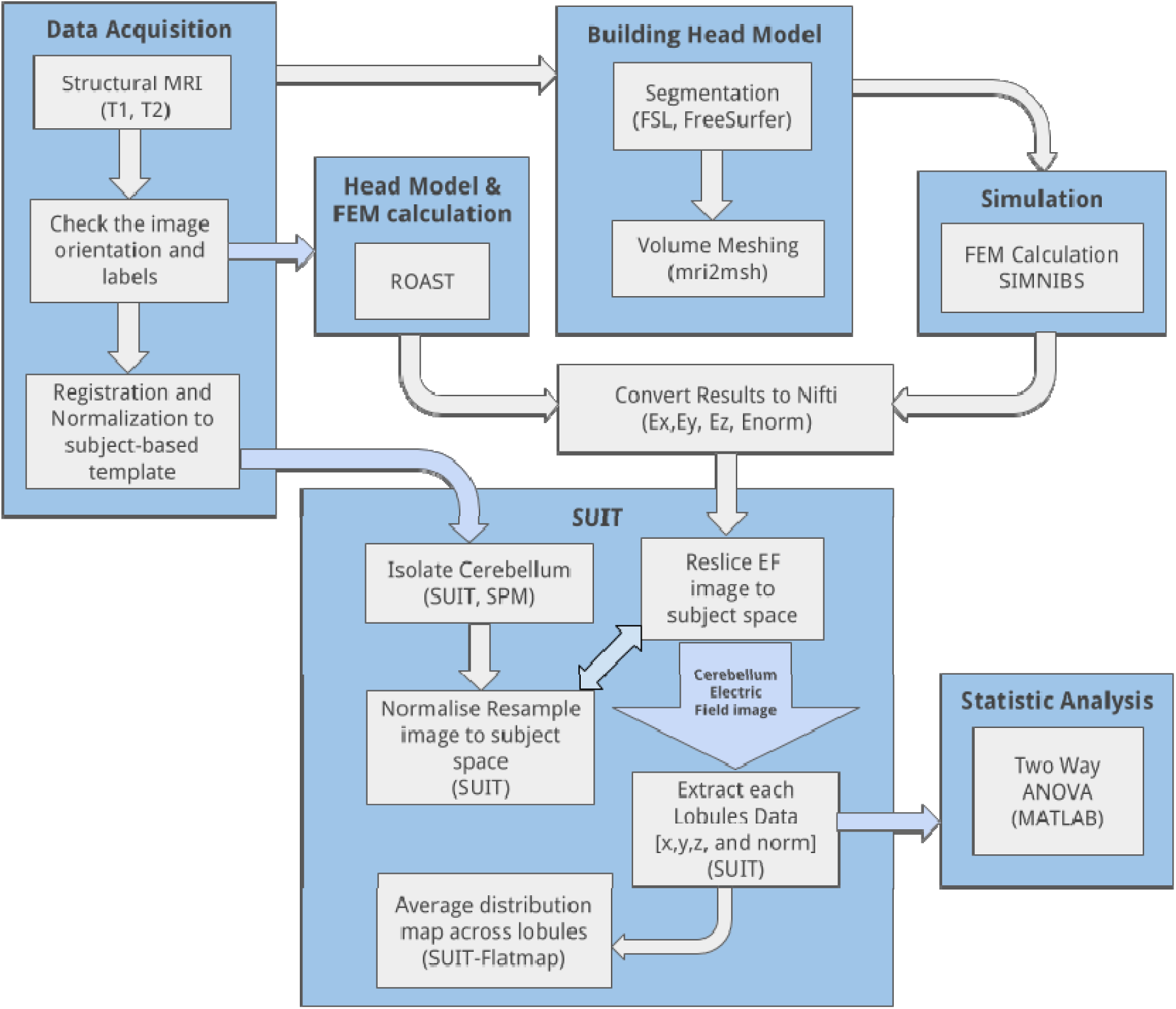
Overall workflow to investigate the electric field distribution across cerebellar lobules during cerebellar transcranial direct current stimulation.

#### 2.1.2. Electrode montages for ctDCS

The electrode positions were defined as follows:

1. **Celnik montage**: 5cm×5cm anode was placed over the right cerebellum, 1 cm below and 3 cm lateral to the inion (Iz, 10/10 EEG system), and the 5cm×5cm cathode was placed over the right buccinator muscle for ctDCS with 2mA direct current.
2. **Manto montage**: 5cm×5cm anode was placed over the right cerebellum, 1 cm below and 3 cm lateral to the inion (Iz, 10/10 EEG system) (Grimaldi and Manto, 2013), and the 5cm×5cm cathode was placed on the contralateral supraorbital area (FP2, 10/10 EEG system) for ctDCS with 2mA direct current.
3. **HD-ctDCS 4×1 montage**: 3.14cm^2^ anode was placed above the cerebellum 10% below Oz (10/10 EEG system) in the midline (Doppelmayr et al., 2016), and four 3.14cm^2^ cathodes were placed at Oz, O2, P8, and PO8 (10/10 EEG system) for ctDCS with 1mA direct current. We investigated the lobular electric field of ctDCS due to the three electrode montages given above using the subject-specific head model as well as the Colin27 head model (Holmes et al., 1998), (Guhathakurta and Dutta, 2016).

#### 2.1.3. Finite element analysis of ctDCS using SimNIBS

Finite element analysis (FEA) was conducted on the subject-specific head model as well as the Colin27 average head model (Holmes et al., 1998), (Guhathakurta and Dutta, 2016) to estimate the ctDCS induced electric field in the brain tissues. The ctDCS was delivered using two 5cmX5cm electrodes and a direct current of 2 mA. In all the simulations, the voxel size was 1mm^3^. The anode and the cathode injected the specified amount of current (source) in the volume conductor, i.e., the head model. The electrodes were modeled as saline-soaked sponge placed at a given scalp location using 10/10 EEG system (Giacometti et al., 2014). We analyzed the head-model for electric field distribution using the SimNIBS pipeline (Windhoff et al., 2013). Following SimNIBS FEA, we used SUIT to isolate the cerebellum in SPM (http://www.fil.ion.ucl.ac.uk/spm/) package in Matlab (The Mathworks Inc., USA). Subject’s T1 images were reoriented into LPI (Neurological) orientation. The isolation map was manually verified in an image viewer (MRIcron). After the isolation, the cerebellum was normalized to the SUIT atlas template using the cropped image and the isolation map. A nonlinear deformation map to the SUIT template is the result of the normalization step. After the normalization, we could either resample the image into SUIT space or into the subject space. The latter was chosen for our subject-specific analysis to resample the probabilistic atlas of the cerebellum into the space of the individual subject. We customized msh2nifti script (https://github.com/ncullen93/mesh2nifti) to save the electric field distribution in the three direction – Ex, Ey, and Ez – from SimNIBS FEA results, as shown by the head model in Figure 2. The msh2nifti script created NIfTI (Neuroimaging Informatics Technology Initiative) images of the electric field distribution that were resliced using the individual mask and deformation matrix (found in the previous step – Figure 1) using the SUIT toolbox to extract the cerebellar regions (or, lobules). The post-processing of the electric field distribution over the tetrahedral volume mesh and its visualization was performed in Gmsh (Geuzaine and Remacle, 2009). The volume of the cerebellar lobules, defined by the SUIT atlas (Diedrichsen, 2006), was used for the extraction of the lobular electric field distribution in Matlab (The Mathworks Inc., USA). To visualise electric field distribution in cerebellar lobules, the flatmap script in SUIT toolbox was used in Matlab (The Mathworks Inc., USA), which provided a flat representation of the cerebellum after volume-based normalization as described by Diedrichsen (Diedrichsen, 2006).

**Figure 2.**
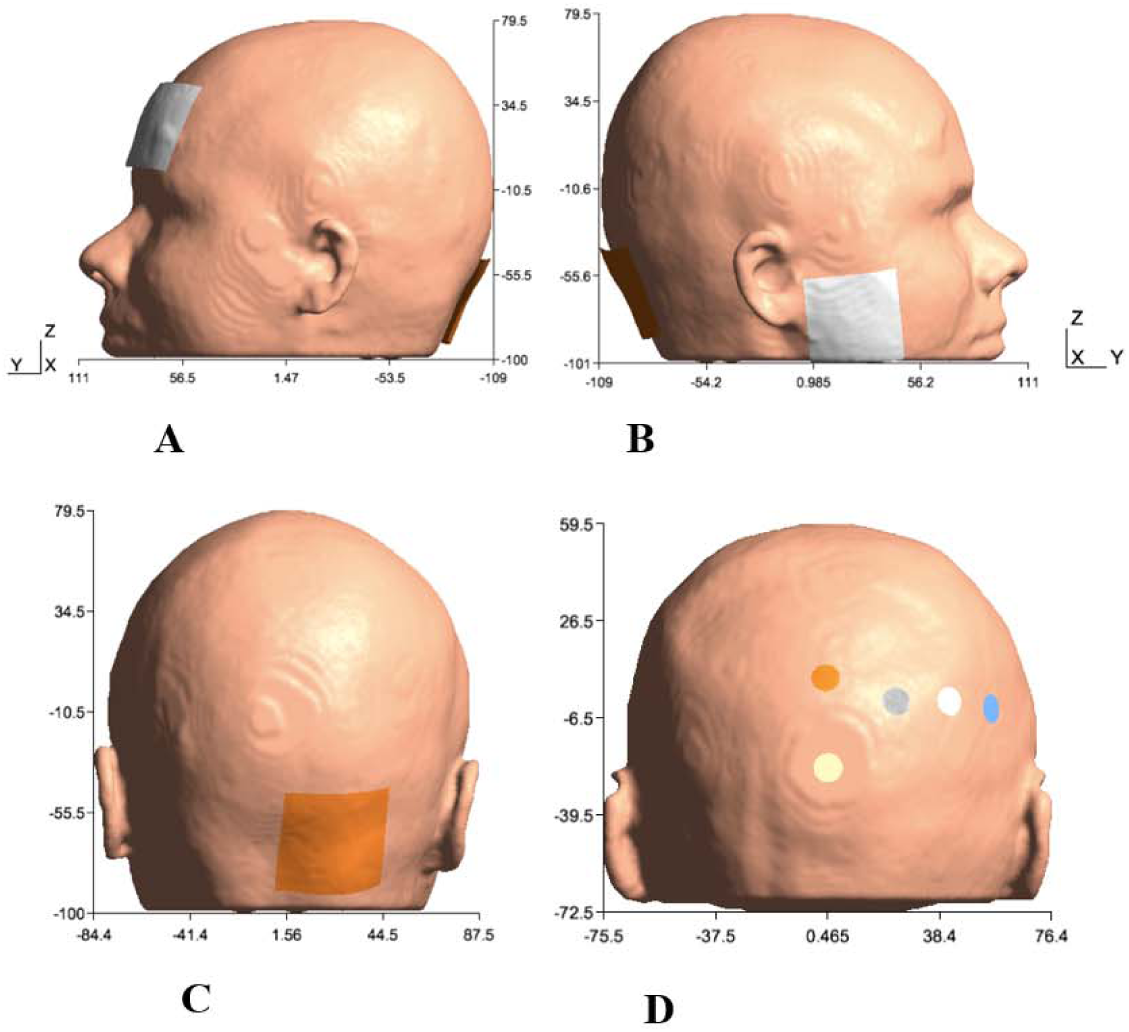
Electrode configurations: A) Manto montage: anode was placed over the right cerebellum, 1 cm below and 3 cm lateral to the inion (Iz, 10/10 EEG system), and the cathode was placed on the contralateral supraorbital area (FP2, 10/10 EEG system), B) Celnik: anode was placed over the right cerebellum, 1 cm below and 3 cm lateral to the inion (Iz, 10/10 EEG system) and cathode was placed over the right buccinator muscle, C) Anode was placed over the right cerebellum, 1 cm below and 3 cm lateral to the inion (Iz, 10/10 EEG system) for Manto and Celnik Montages, and D) HD-tDCS: anode was placed above the cerebellum 10% below Oz (10/10 EEG system) in the midline, and four cathodes were placed at Oz, O2, P8, and PO8 (10/10 EEG system).

For group analysis of the lobular electric field distribution, an averaging of the SUIT flatmap across subjects is possible. Here, the electric field distribution across lobules is important to determine the focality of different ctDCS montages. We performed the analysis of variance (ANOVA) of the lobular electric field distribution using ANOVA to investigate the factors of interest – lobules (28 from SUIT), montages (Celnik, Manto, HD-ctDCS), head model (Colin27, subject-specific), and their interactions. Post-hoc multiple comparison test was conducted using Bonferroni critical values. Also, in the Generalized Linear Model (GLM), the proportion of the total variability in the dependent variable that is accounted for by the variation in the independent variable was found using the eta-squared effect size measure (Lakens, 2013).

#### 2.1.4. Finite element analysis of ctDCS using ROAST

We used another FEA pipeline called Realistic volumetric Approach to Simulate Transcranial Electric Stimulation (ROAST) (Huang et al., 2018) to compare the lobular electric field distribution with that from the SimNIBS pipeline (section 2.1.3). We constructed subject-specific head model using the same T1- and T2-weighted MRI from section 2.1.3. The creation of the tetrahedral volume mesh of the head and solving the finite element model were implemented by ROAST. The pipeline is a Matlab script based on three open source software: Statistical Parametric Mapping (SPM) (Penny et al., 2011), Iso2mesh (Fang and Boas, 2009), and getDP (Dular et al., 1998). ROAST provided results for electric field distribution as NIFTI images which were processed in our pipeline to isolate the cerebellum and analysis of the lobular electric field distribution, as described in section 2.1.3.

### 2.2. Experimental Data from Healthy Human Study

In our healthy human study (experimental results published in conference proceedings (Abadi and Dutta, 2017)), fifteen healthy volunteers participated in accordance with the Declaration of Helsinki. Ethics approval was obtained at the University Medical Center, Goettingen, Germany. In this study, two-electrode ctDCS montages were investigated for the application of anodal transcranial direct current stimulation over the cerebellar hemisphere during visuomotor learning of myoelectric visual pursuit using the electromyogram (EMG) from ipsilateral gastrocnemius (GAS) muscle. This study was conducted to investigate the effects of 15min of anodal ctDCS (current density=0.526A/m^2^; electrode size 5cm×5cm) using Celnik and Manto montage on the response time (RT) and root mean square error (RMSE) during isometric contraction of the dominant GAS for myoelectric visual pursuit, i.e., ‘ballistic EMG control’ (Dutta et al., 2014) (see (Abadi and Dutta, 2017) for further details). The EMG RT was computed offline as the duration from the instant of visual cursor target cue to the instant when the rectified EMG in a sliding window of 500ms from the muscle jumped by more than three times of the standard deviation of the resting value. The response accuracy was computed as RMSE between the EMG driven cursor and the cursor target signals during cue presentation. 95% confidence intervals for the parameters were compared for overlap between post-intervention and baseline based on Student’s t-distribution.

## 3. Results

### 3.1.1. Finite element analysis of ctDCS using SimNIBS

Figure 3 shows a higher average electric field magnitude at the right cerebellar hemisphere than the left cerebellar hemisphere since the anode was placed over the right cerebellum (3 cm lateral to the inion) in both the Celnik and the Manto montages. FEA using SimNIBS showed that the ctDCS electric field magnitude for both the Celnik and the Manto montages can spread to neighboring structures, e.g., the right temporal lobe for the Celnik montage and the left prefrontal cortex for the Manto montage, as shown in Figure 3A and 3B. Moreover, ctDCS electric field magnitude for the HD-ctDCS montage by Doppelmayr and colleagues (Doppelmayr et al., 2016) can spread to the occipital lobe, as shown in Figure 3C. Here, the HD-ctDCS anode was placed 10% below the inion, but the cathodes were located at Oz, O2, P8, and PO8 which are partly on the right occipital lobe. Furthermore, the spread to the occipital lobe can be facilitated by the highly conductive CSF.

**Figure 3.**
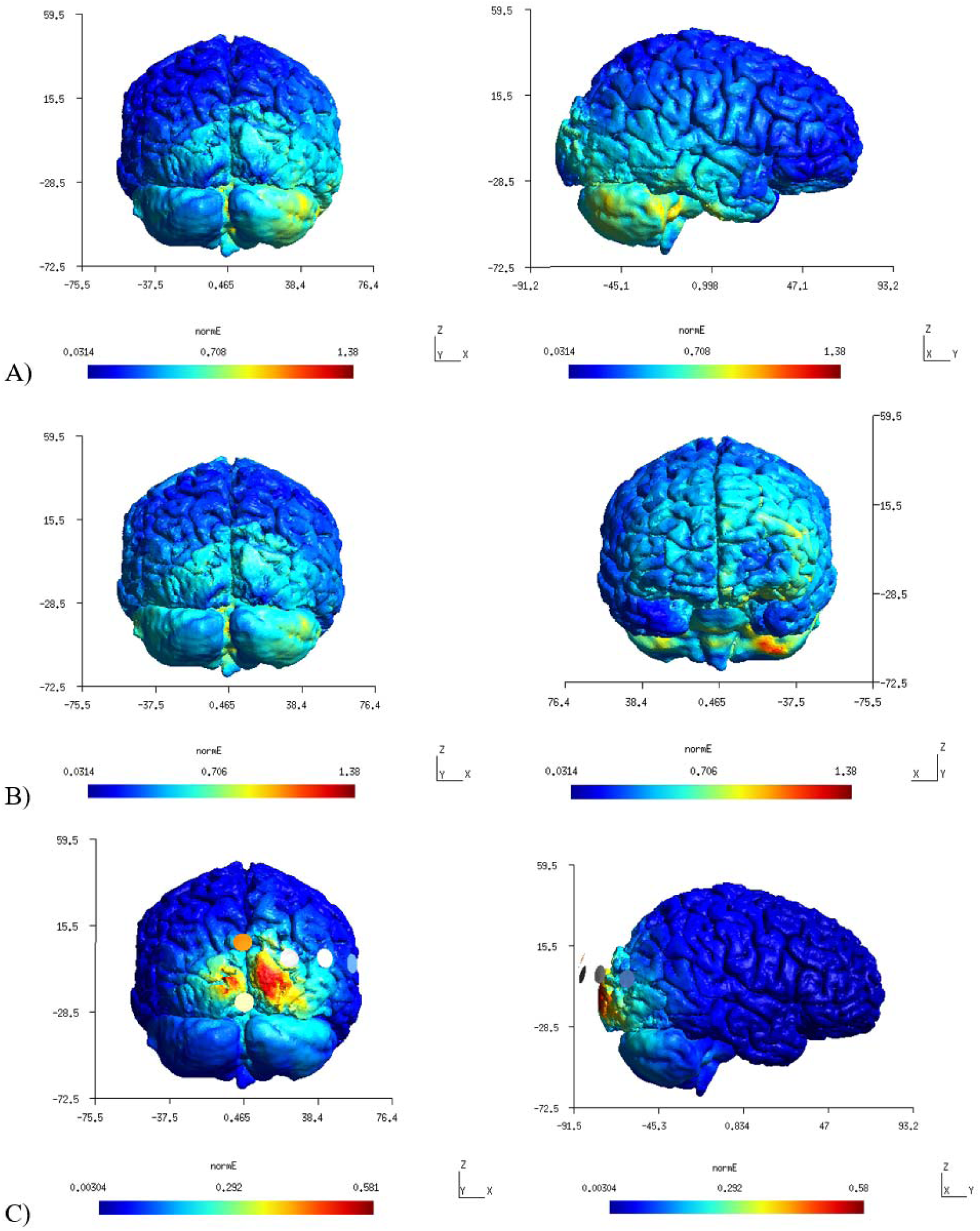
A) First-row panels (color scale: 0.0314 to 1.38 V/m) – electric field distribution for Celnik montage which was found confined to the right hemisphere. B) Second-row panel (color scale: 0.0314 to 1.38 V/m) – electric field distribution for the Manto montage at the right cerebellum and left prefrontal cortex. C) Third-row panels (color scale: 0.00304 to 0.581 V/m) – electric field distribution for the HD-tDCS montage.

We further analyzed the SimNIBS FEA results using our SUIT-based computational pipeline to compute lobular electric field distribution. The SUIT flatmap results for Ex, Ey, Ez, and Enorm (electric field magnitude or strength) are shown in Figure 4A. Ey, which is approximately normal to the surface at the anode, has the highest strength (highest 0.6V/m) while Ex and Ez have comparable strength (highest 0.3V/m). Celnik and Manto montages primarily affected the posterior and the inferior parts of the right cerebellum, as shown in Figure 4A by the SUIT flatmap Enorm results. However, Manto montage had a more bilateral effects on the cerebellar hemispheres than the Celnik montage based on the electric field strength (Enorm). When compared with the Colin27 head model, our subject-specific head model resulted in an overall lower magnitude electric field distribution. Moreover, one can visualize the subject-specific differences in the lobular electric field distribution with the Colin27 head model which shows the importance of subject-specific optimization of the electrode montage. With the HD-ctDCS montage, the electric field strength (Enorm) was found to be more focal at Crus I, Crus II of the targeted right cerebellum, as shown in Figure 4B, although the magnitude was much lower due to a smaller (1mA) injected direct current at anode. The current density at the electrode skin interface was only 0.08mA/cm^2^ for the Celnik and the Manto montages but much higher at 0.32mA/cm^2^ for the HD-ctDCS anode.

**Figure 4.**
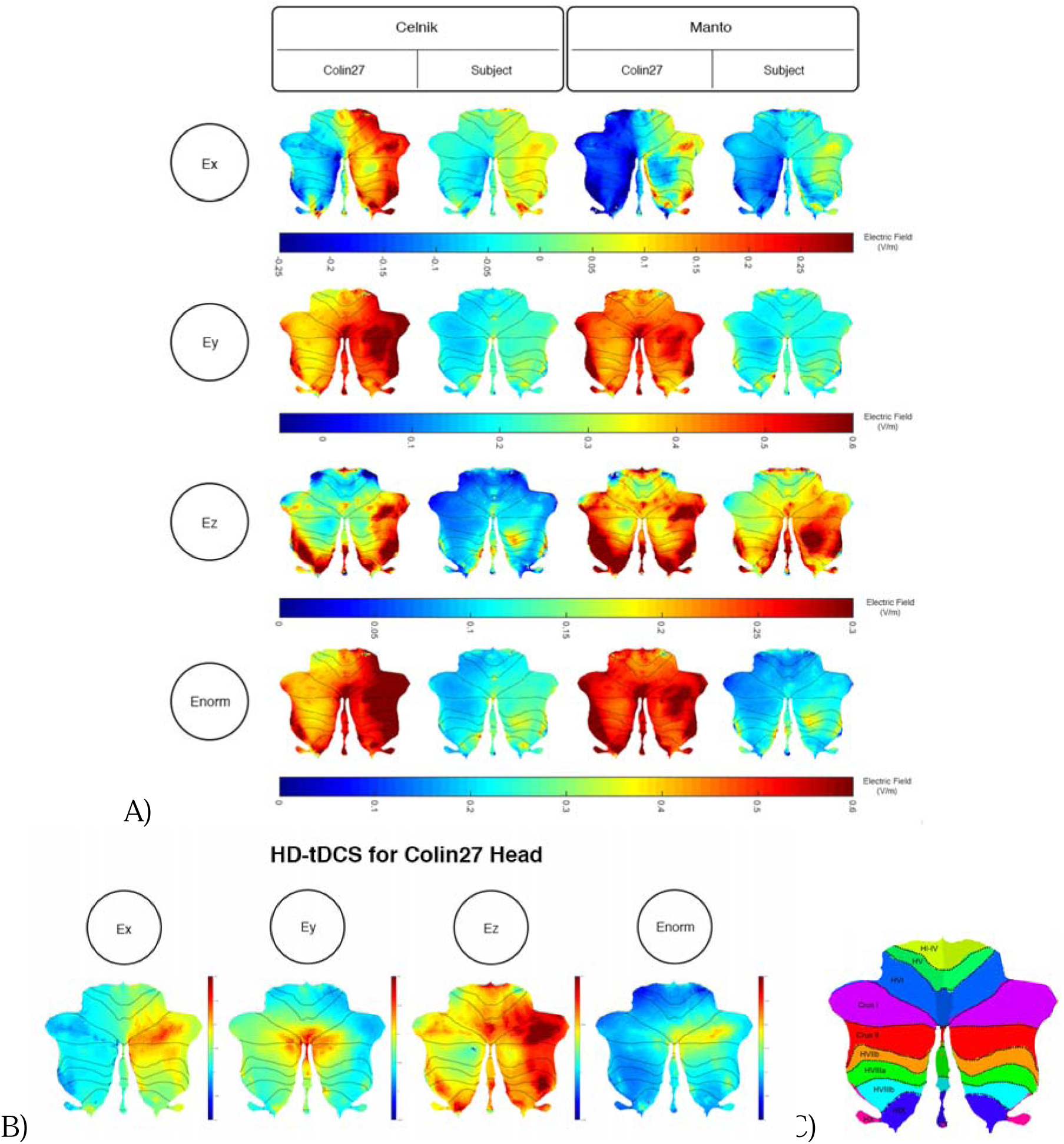
A) Comparison between the SimNIBS outcomes for Celnik and Manto Montages on Colin27 and subject-specific head model. Electric field distribution (Ex, Ey, Ez, and Enorm) of Celnik and Manto montages for Colin27 Head and Subject-specific head model were visualized in SUIT toolbox using flatmap. First row: Color Scale of –0.25 to 0.3 V/m – Electric field distribution (Ex); second row: Color Scale of −0.05 to 0.6 V/m – Electric field distribution (Ey); third row: Color Scale of 0 to 0.3 V/m – Electric field distribution (Ez); fourth row: Color Scale of 0 to 0.6 V/m – Electric field distribution (Enorm). **B) Electric field distribution of HD-tDCS montage for Colin27 head model**: Ex (color scale: −0.02 to 0.06 V/m), Ey (color scale: 0 to 0.18 V/m), Ez (color scale: 0 to 0.5 V/m), and Enorm (color scale: 0 to 0.15 V/m). **C) SUIT lobules** (Diedrichsen et al., 2009)

The analysis of variance (ANOVA) results were evaluated for statistical significance using the eta-squared effect size measure. In order to investigate the effect of head model on the electric field strength (Enorm), we computed a three-way ANOVA, and found that the eta-squared effect size was 0.05 for lobule, 0.00 for montage, 0.04 for head model, 0.01 for lobule*montage interaction, 0.01 for lobule* head model interaction, and 0.00 for montage*head model interaction in case of Enorm. The lobule*head model interaction for the electric field strength (Enorm) is shown in Figure 5, which shows that the magnitudes are different even though the overall distribution is similar. If we consider only the subject-specific head model then the eta-squared effect size measure from two-way ANOVA results was 0.03 for lobule, 0.05 for montage, and 0.02 for interaction in case of Enorm; 0.38 for lobule, 0.02 for montage, and 0.07 for interaction in case of Ex; 0.03 for lobule, 0.05 for montage, and 0.02 for interaction in case of Ey; 0.09 for lobule, 0.04 for montage, and 0.04 for interaction in case of Ez. Here, the effect sizes are mostly small except for lobule*montage interaction for Ex, and lobule for Ez which are moderate.

**Figure 5.**
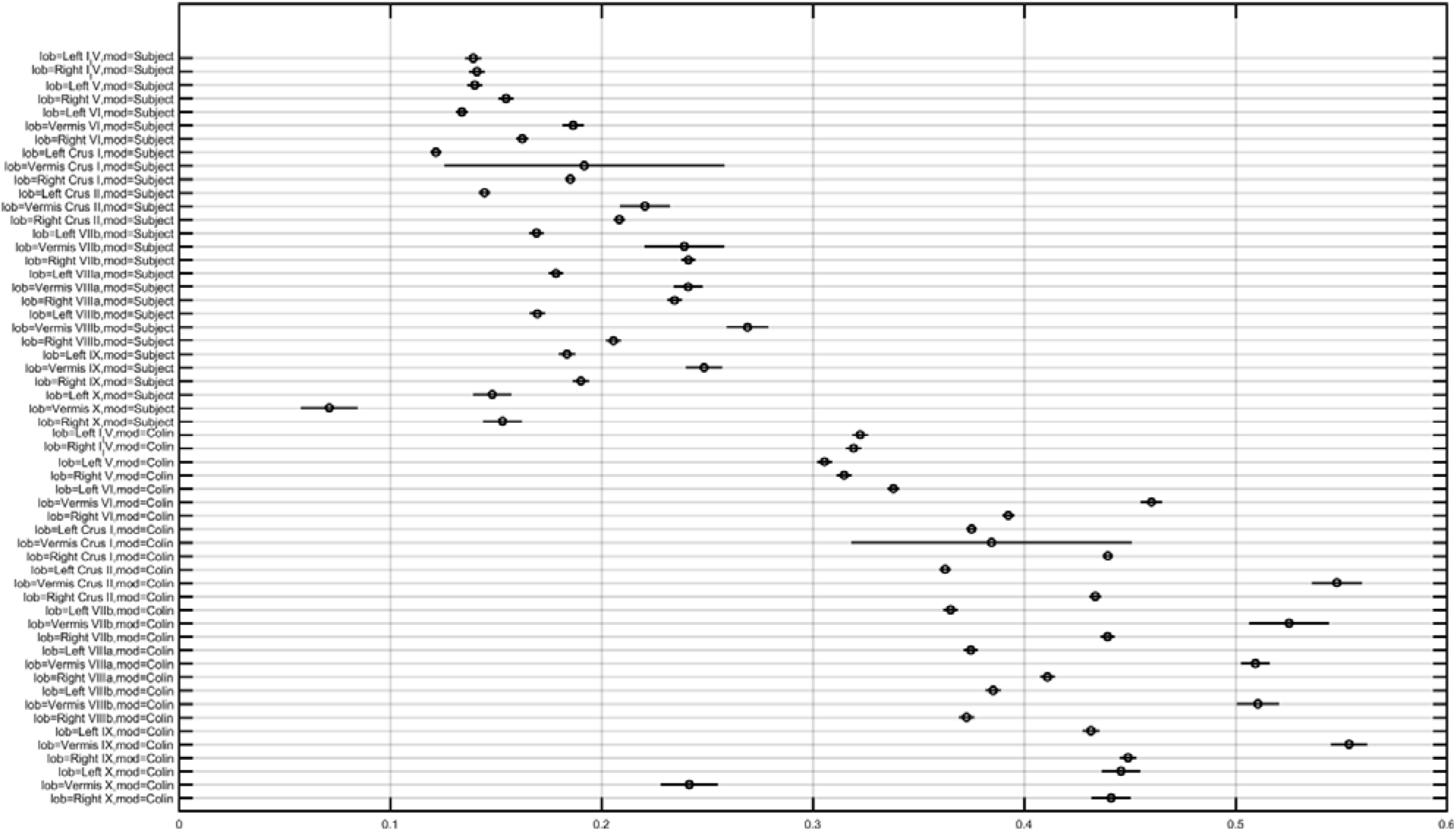
Multiple comparison results for lobule*head model interaction for electric field strength (Enorm) (X-axis in V/m).

Two-way ANOVA of the lobular electric field strength (Enorm) with montages and the post-hoc multiple comparisons of the means (95% significance) with Bonferroni critical values showed that the Celnik and Manto montages primarily affected the lobules Crus II, VIIb, VIII, IX of the targeted right cerebellar hemisphere – see Figure 1A in the supplementary material. However, Manto montage was found to have a more bilateral effect on the electric field strength (Enorm) which is also visible in the Figure 4A. Enorm for HD-ctDCS primarily affected the lobules Crus I, Crus II, VIIb of the targeted right cerebellar hemisphere – see Figure 1A in the supplementary material. Post-hoc multiple comparison of the means (95% significance) of the Ex, Ey, and Ez are shown in the Figures 1B, 1C, and 1D in the supplementary material. An interesting finding is the opposite direction of the Ex electric field in most cerebellar lobules in the Manto montage when compared to the HD-ctDCS montage, and the Celnik montage is in the middle having both directions. Post-hoc multiple comparisons on Ex, Ey, Ez also revealed that Ey lobular distribution was different for the HD-ctDCS montage when compared to Manto and Celnik montages (as shown in Figures 1C in the supplementary material) where Ey in HD-ctDCS primarily targeted the vermis region. The average electric field strength (Enorm) across different lobules for Celnik montage is listed in Table 2 in the supplementary material. In the supplementary material, Table 3 lists for Manto montage and Table 4 lists the same for the HD-ctDCS montage.

### 3.1.2. Comparison of finite element analysis using SimNIBS and ROAST

SimNIBS runs much slower (~8-10hrs) when compared to ROAST (15-30mins) that leverages the volumetric segmentation from SPM (Huang et al., 2017). Huang and colleagues (Huang et al., 2017) have shown a high deviation of SimNIBS-generated electric field compared to SPM-generated result in ROAST which was also found in our the lobular electric field distribution results – see Figure 6. Huang and colleagues (Huang et al., 2017) postulated that the source of this difference comes mainly from the two different segmentation approaches, and described the volumetric approach of segmentation in the ROAST pipeline being more realistic of the anatomy when compared to the surface-based segmentation in SimNIBS. We found that the components with intersecting surfaces (e.g., gray matter and cerebellum) was better capture by ROAST which led to a better estimate of the bilateral electric field Ez and Enorm in the Manto montage – see Figure 6. This indicates a genuine difference in these two categories of modeling methods where ROAST performed better (also highlighted by (Huang et al., 2017)). The limitation with SimNIBS is the difficulty in capturing the fine details of the cerebellum which is important in computing the lobular electric field distribution.

**Figure 6.**
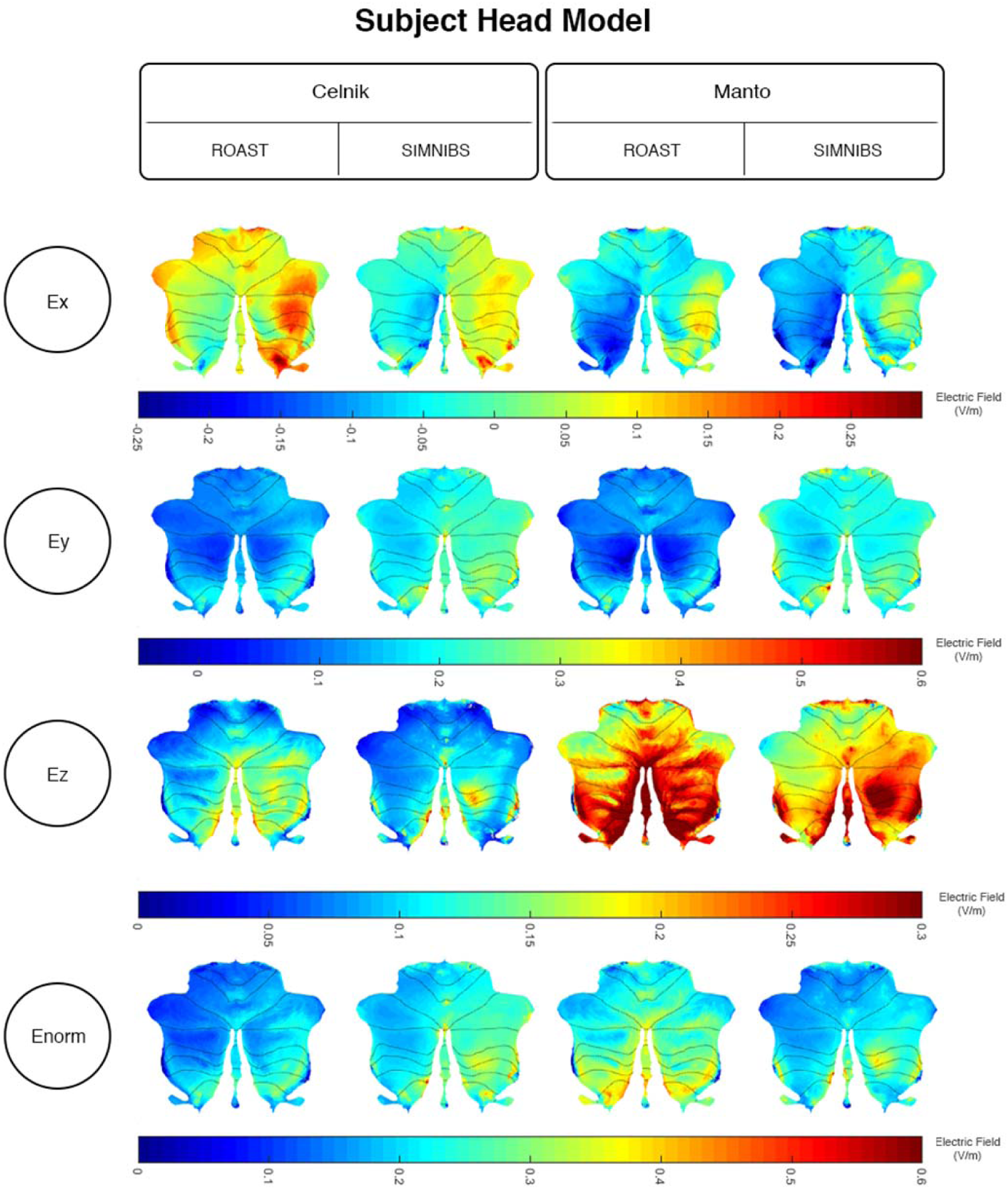
Comparison of the SimNIBS and ROAST outcomes for Celnik and Manto Montages for the subject-specific head model. Electric field distribution (Ex, Ey, Ez, and Enorm) of Celnik and Manto montages for Subject-specific head model were visualized in SUIT toolbox using flatmap. First row: Color Scale of –0.25 to 0.3 V/m – Electric field distribution (Ex); second row: Color Scale of −0.05 to 0.6 V/m – Electric field distribution (Ey); third row: Color Scale of 0 to 0.3 V/m – Electric field distribution (Ez); fourth row: Color Scale of 0 to 0.6 V/m – Electric field

### 3.1.3. Analysis of the experimental results using our computational pipeline

The computational SUIT-based analysis presented in this technology report was used to investigate healthy human anodal ctDCS results during a VMT performance (Foerster et al., 2015). Our prior experimental results (Abadi and Dutta, 2017) showed that Manto montage resulted in a statistically significant (p<0.05) decrease in RT post-intervention than baseline when compared to the Celnik montage while Celnik montage resulted in a statistically significant (p<0.05) decrease in RMSE post-intervention than baseline when compared to the Manto montage. In fact, cerebellar tDCS using Celnik montage has shown to affect the adaptation rate of spatial but not temporal elements of walking (Jayaram et al., 2012) which was postulated to be related to electric field effects on different cerebellar regions, e.g., vermis (for spatial) versus adjacent hemispheres (for temporal elements) (Jahn et al., 2004). Indeed, we found in the analysis using our computational pipeline that Celnik montage has more unilateral effect of the electric field strength (Enorm) on the cerebellar hemispheres including vermis, as shown in Figure 6, when compared to Manto montage that has bilateral effect of the electric field strength (Enorm). Such bilateral effect of the electric field strength (Enorm) may be responsible for a statistically significant (p<0.05) decrease in RT post-intervention than baseline, i.e., temporal aspects (Jahn et al., 2004) affected by the Manto montage. Here, Celnik montage resulted in a statistically significant (p<0.05) decrease in RMSE post-intervention than baseline which is a spatial aspect of the target pursuit during VMT (Abadi and Dutta, 2017). The lobular electric field distribution shown in Figure 6 indicated electric field primarily in the posterior-anterior (Y) and inferior-superior (Z) directions for both the Celnik and the Manto montages. However, the electric field in the mediolateral (X) direction changed direction across hemispheres in the Manto montage, going from a minimum of −0.25V/m in the left cerebellum to a maximum of 0.3V/m in the right cerebellum, which is in contrast to that in the Celnik montage that stays mostly positive from 0 – 0.3V/m. Here, the electric field strength (Enorm) primarily affected the lobules Crus II, VIIb, VIII, IX (compare with Figure 4C) of the targeted cerebellar hemisphere.

## 4. Discussion

Our freely available computational modeling approach to analzye subject-specific lobular electric field distribution during ctDCS provided an insight into healthy human anodal ctDCS results during a VMT performance (Foerster et al., 2015),(Abadi and Dutta, 2017). Bilateral effect of the electric field strength (Enorm) due to Manto montage is postulated to be responsible for a statistically significant (p<0.05) decrease in RT post-intervention than baseline, i.e., temporal aspects (Jahn et al., 2004), while unilateral electric field distribution due to Celnik montage affected the spatial aspect of the target pursuit during VMT (Abadi and Dutta, 2017) and resulted in a statistically significant (p<0.05) decrease in RMSE post-intervention than baseline. The Celnik and Manto montages in the subject-specific head model affected primarily the lobules Crus II, VIIb, VIII, IX of the targeted cerebellar right hemisphere as shown in Figure 6. The ctDCS effects on RMSE by Celnik montage may be the result of motor adaptation based on the tDCS-modulation of the synaptic activity between parallel fiber and the dendritic tree of the Purkinje cells (i.e., the matrix memory) while the decrease in RT to unanticipated visual cue by Manto montage may be the result of ctDCS-enhanced responsiveness of the Purkinje cells (the sole output of the cerebellar cortex) – see Ez in Figure 7 (for the cerebellar lobules related to lower limb function). Here, cerebellum serves two purposes according to the computational model, i.e., Purkinje cells provide both the internal prediction and sensory feedback signals for motor control (Popa et al., 2016).

**Figure 7.**
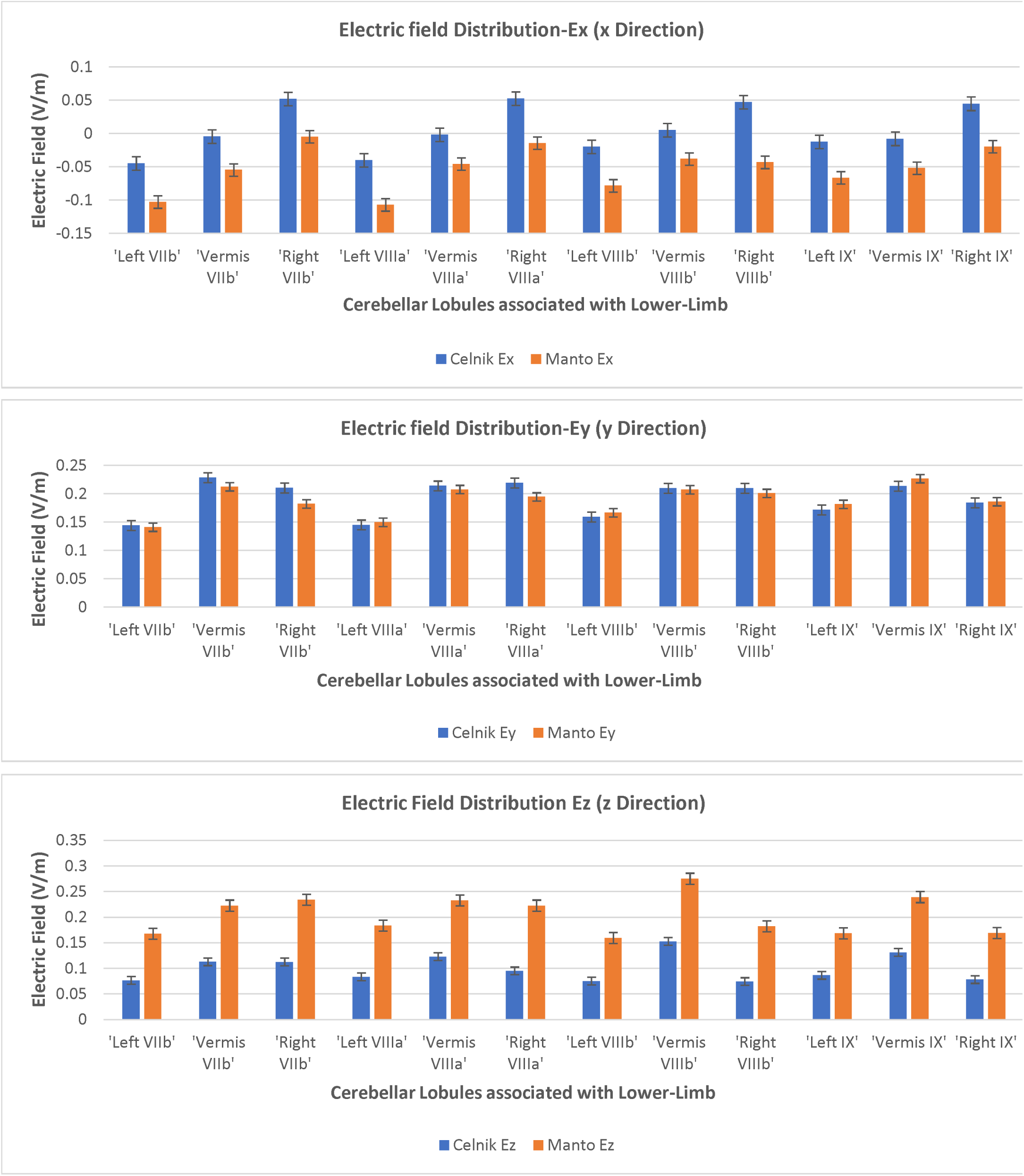
Electric Field Distribution in X, Y, Z directions for the lobules associated with lower-limb function.

Figure 7 shows the results for the cerebellar lobules related to lower limb function. This is relevant for posture and gait which are sensorimotor actions that involve peripheral, spinal, and supraspinal structures (Jahn et al., 2004). Impaired standing balance, and abnormal eye movements are common problems in persons with multiple sclerosis (pwMS) which may occur from brainstem and cerebellum lesions that disrupt the integration of signals from the visual, somatosensory, and vestibular systems (Bennett and Leavitt, 2018). Moreover, movement inefficiency and postural control impairment in MS may lead to falls and fatigue in pwMS. Therefore, locomotor rehabilitation needs to address efficient sensory motor integration with balance and eye movement exercises (BEEMS) (Hebert et al., 2018) while the subject processes multi-sensory information (Bennett and Leavitt, 2018). Here, the mechanisms underlying BEEMS effects on conduction delays (ostensibly as a result of a demyelinating lesion) during dynamic balance in pwMS needs further investigation. During BEEMS, one type of sensorimotor learning may involve building a specific internal model of the environment for feedforward prediction (Thoroughman and Shadmehr, 1999) to overcome time delays associated with feedback control (Wolpert et al., 1998). Cerebellar architecture has been found to support the computations required by such a feedforward model from animal studies as well as patients with cerebellar dysfunction (Ebner, 2013) since the Purkinje cell firing has several of the characteristics of a forward internal model (Ebner, 2013) (Thoroughman and Shadmehr, 2000). During BEEMS, the neural circuits of the brain and spinal cord will respond to intrinsic (volitional drive during exercises) and extrinsic stimuli by reorganizing their structure, function, and connections, which is called neuroplasticity (Cramer et al., 2011). Here, the strength of cerebellar-to-cerebral pathways for a upper limb and lower limb muscle may reflect novel somatotopy of cerebellum-dependent motor adaptation (Spampinato et al., 2017) during BEEMS.

Marr-Albus-Ito hypothesis combined with the computational model of the cerebellum as an internal model (Popa et al., 2016) can provide insights into the relevance of the electric field direction in the cerebellar cortex. Preliminary human studies hypothesized that ctDCS effects were due to polarizing effects on Purkinje cells (Galea et al., 2009) however a recent animal study (Sánchez-León et al., 2017) reflected the importance of the synaptic activity between parallel fiber and the dendritic tree of the Purkinje cells. This aligns well with the Marr-Albus-Ito hypothesis that the strength of the parallel fiber synapse when the climbing fiber fires will determine the Purkinje cell’s firing. Here, due to the primarily orthogonal orientation of the parallel fibers concerning the climbing fiber and Purkinje cells in the cerebellar cortex, the ctDCS electric field direction is postulated to be relevant. It is noteworthy that although Galea and colleagues (Galea et al., 2009) initially replicated the effect of ctDCS on visuomotor adaptation, this was not maintained during a large range of varying task parameters questioning the validity of using ctDCS within a clinical context (Jalali et al., 2017). We postulate that subject-specific ctDCS electric field orientation within the cerebellar lobules needs to be investigated within a clinical study based on realistic head modeling where our computational pipeline will be useful. When comparing the Celnik and Manto montages, we found that the electric field in the mediolateral (X) direction might affect the parallel fibers differently between Celnik and Manto montages due to difference in the Ex direction, as shown in Figure 6. The top panel of the Figure 7 shows that the electric field in the mediolateral (X) direction is all negative for Manto montage when compared to Celnik montage for the targeted right hemisphere. Here, the direction of electric field vector requires investigation using multi-scale modeling (Seo and Jun, 2017) vis-à-vis Purkinje cell, climbing fiber, and parallel fiber orientations, which is our future work. Also, HD-ctDCS montage primarily affected the lobules Crus I, Crus II, VIIb of the targeted cerebellar hemisphere that are linked to cognitive impairments (Stoodley and Schmahmann, 2010). Therefore, HD-tDCS montage presented by Doppelmayr and colleagues (Doppelmayr et al., 2016) may be relevant for cognitive rehabilitation.

One limitation of this technology report is the lack of neurophysiological testing. For example, cerebellar brain inhibition (CBI) is a physiological parameter of the connectivity strength between the cerebellum and the primary motor cortex (M1) that can be identified using transcranial magnetic stimulation (TMS) (Fernandez et al., 2018). Moreover, one may need to use in vivo intracranial cerebellar recordings in humans for experimental validation (Huang et al., 2017). Another limitation is an uncertain (anisotropic) conductivity profile for the cerebellum that can have a substantial influence on the prediction of optimal stimulation protocols, e.g., (Schmidt et al., 2015). These needs to be addressed in the future based on neurophysiological testing. Indeed, individualized protocol for ctDCS that is verified with neurophysiological testing is necessary to reduce inter-individual variability (Iodice et al., 2017). Therefore, our freely available computational pipeline for cerebellum that are easily accessible worldwide is crucial to facilitate clinical translation of ctDCS for neurorehabilitation, e.g. using VMT. Here, system analysis using an error clamp design of the VMT (Kha et al., 2018) may further elucidate the behavioral mechanism of ctDCS where no visual feedback is presented after motor adaptation during ‘error clamp’ trials (Vaswani and Shadmehr, 2013). It has been postulated that motor memories show little decay in the absence of error if the brain is prevented from detecting a change in task conditions (Vaswani and Shadmehr, 2013). Therefore, during ‘error clamp’ trials, Celnik montage should have little effect on both RMSE and RT (also the case with visual feedback) while Manto montage is postulated to have a significant effect on RT (same with no visual feedback) and little effect on RMSE (also the case with visual feedback). This will indicate that Manto montage’s RT effects are solely due to sensory feedback signals from the Purkinje cells. We have found RT effect of primary motor cortex (M1) tDCS that changed the input-output function of the pyramidal cells (Lafon et al., 2017) leading to response time improvement post-tDCS when compared to pre-tDCS baseline performance (Kha et al., 2018). Also, a systematic evaluation of more focal electrode montages, such as high definition (HD) tDCS (Doppelmayr et al., 2016), may elucidate the specificity of the ctDCS effects. HD-ctDCS may reach deeper into the cerebellum while limiting diffusion to neighboring structures(Fiocchi et al., 2017). From the Doppelmayr’s study [17], we expected a more focal electric field distribution in HD-ctDCS montage. However, we found diffusion to neighboring structures, e.g., the occipital lobe, as shown in Figure 4B. Therefore, visual cortex effects of HD-ctDCS need to be delineated in such visuomotor adaptation studies in the future.

The interaction between the shape of the lobule and the electric field gradient determines the variability of the lobular electric field distribution, e.g., in the vermis region of Crus I, that can be found in Figure 4 (Tables 1, 2, 3 in the supplementary material). If the major axis of the geometry is aligned along the major axis of the electric field gradient at the centroid, then we get a large variance. We postulate that not only the mean but also the variability of the electric field distribution at a brain location is important since the tDCS effects are due to electric field ostensibly modulating population rate and spike timing (Reato et al., 2010). Therefore, subject-specific customization of electrode placement need to take into account both the mean and the variance of the electric field distribution that can be affected due to the complexity of cerebellar structure. Our lobular modeling approach in aligning the major axis of the geometry of the brain region with the minor axis of the electric field gradient at the centroid while maximizing the average electric field strength will be relevant for optimization of multi-channel electrode montage (Otal et al., 2016). For example, optimizing ctDCS targeting Crus I, Crus II as an adjuvant treatment for cognitive training (Bernard and Seidler, 2013) may need to reduce the variability in the electric field distribution across the vermis. Here, electrode size is important since it determines the current density at the electrode-skin interface which is primarily responsible for the cutaneous pain sensation (Minhas et al., 2011). Therefore, a relatively higher 2 mA current intensity for Celnik and Manto montages is tolerable at the electrode-skin interface since they use larger (5cm x 5cm) electrodes when compared to HD-ctDCS montage (1cm radius circular electrode). However, the total current injected by the anode primarily determines the electric field strength at the targeted brain region. Here, the cerebellum is located more in-depth from the scalp, and a significant amount of current gets shunted by the cerebrospinal fluid (Rampersad et al., 2014), so we may need more current intensity at the anode that has to be tolerable at the electrode-skin interface. Optimization using multi-anode (1cm radius) tDCS (Otal et al., 2016) instead of single anode in 4×1 HD-ctDCS will distribute the total current across anode and can provide more focal targeting of Crus I, Crus II, VIIb (Stoodley and Schmahmann, 2010) for cognitive training. Moreover, the direction of the electric field vector may be better controlled by current steering using multi-channel montage (Otal et al., 2016), e.g., in aligning the major axis of the electric field gradient with the cerebellar peduncles, which may be relevant for motor neurorehabilitation (Dutta et al., 2014) (Abadi and Dutta, 2017).

### Author Contributions

ZR conducted the computational modeling under the guidance of AD. All the authors have drafted the work and revised it critically and have approved the final version before submission.

### Ethics Approval

For research participation as well as the publication of the case report including the participant's identifiable information, a written informed consent was obtained from the subject at the University at Buffalo prior to presenting this case. Ethics approval for the experimental study on visuo-myoelectric control was obtained at the University Medical Center, Goettingen, Germany.

## Acknowledgement

AD would like to acknowledge the help received from Dr. med. M.A. Nitsche at the University Medical Center, Goettingen, Germany for the data collection during visuo-myoelectric control study.

## Funding

University at Buffalo SUNY funded this computational work.

## Conflict of Interest Statement

The authors declare that they have no conflict of interest.

