## Supplemental Data for "A computational pipeline to determine lobular electric field distribution during cerebellar transcranial direct current stimulation"

**Supplementary Figure 1**

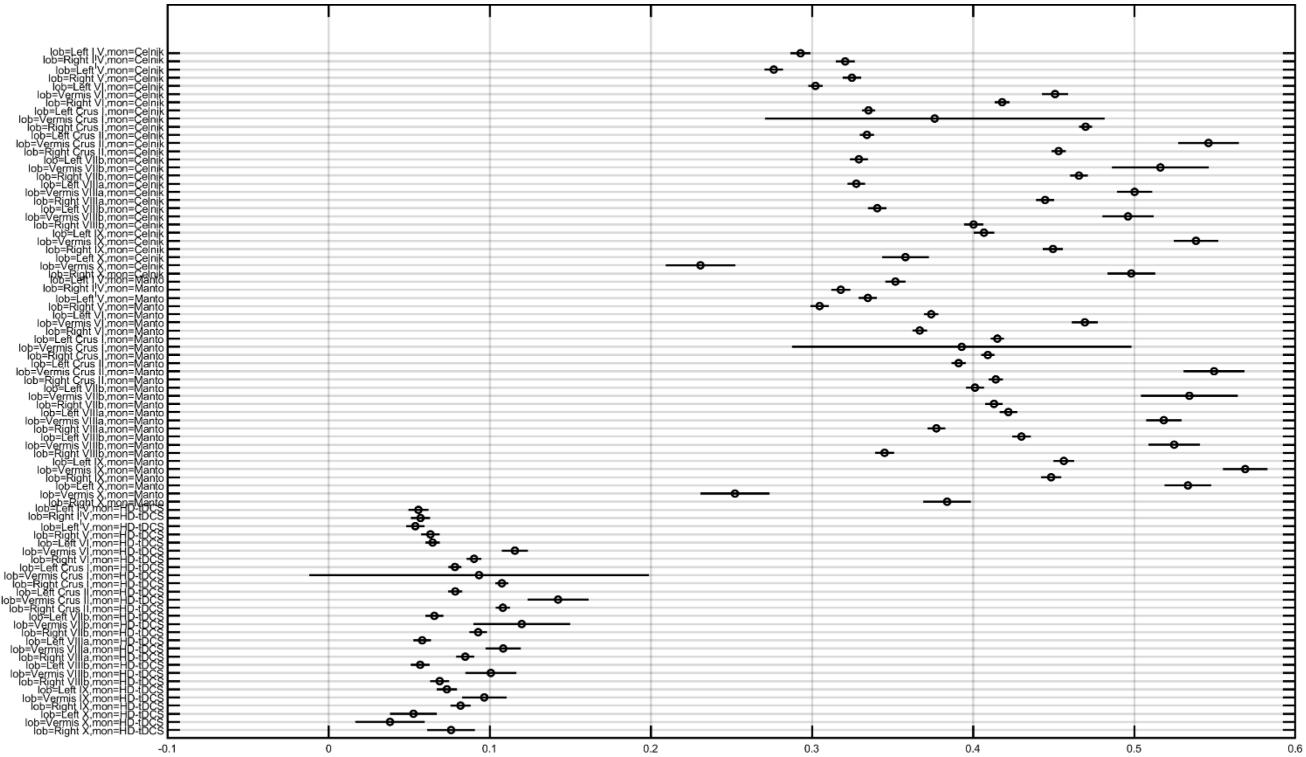


A)

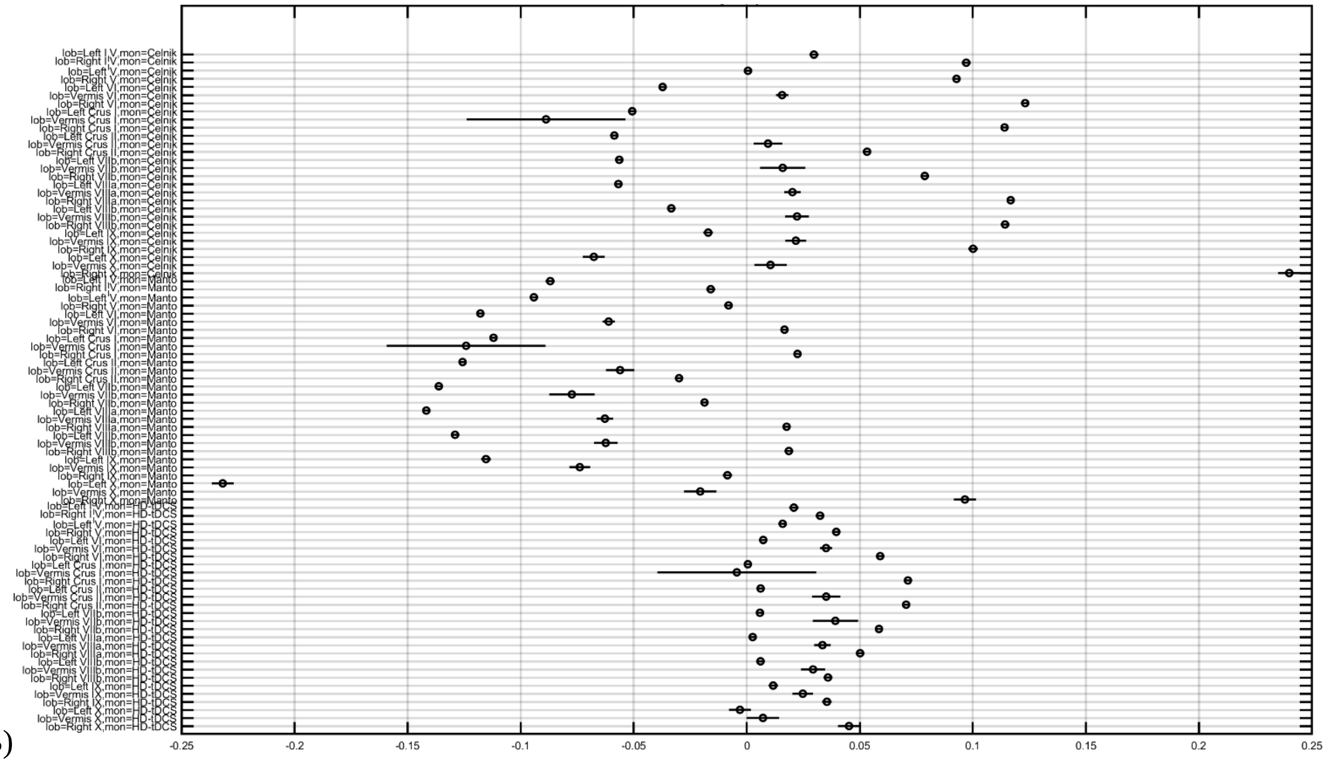


B)

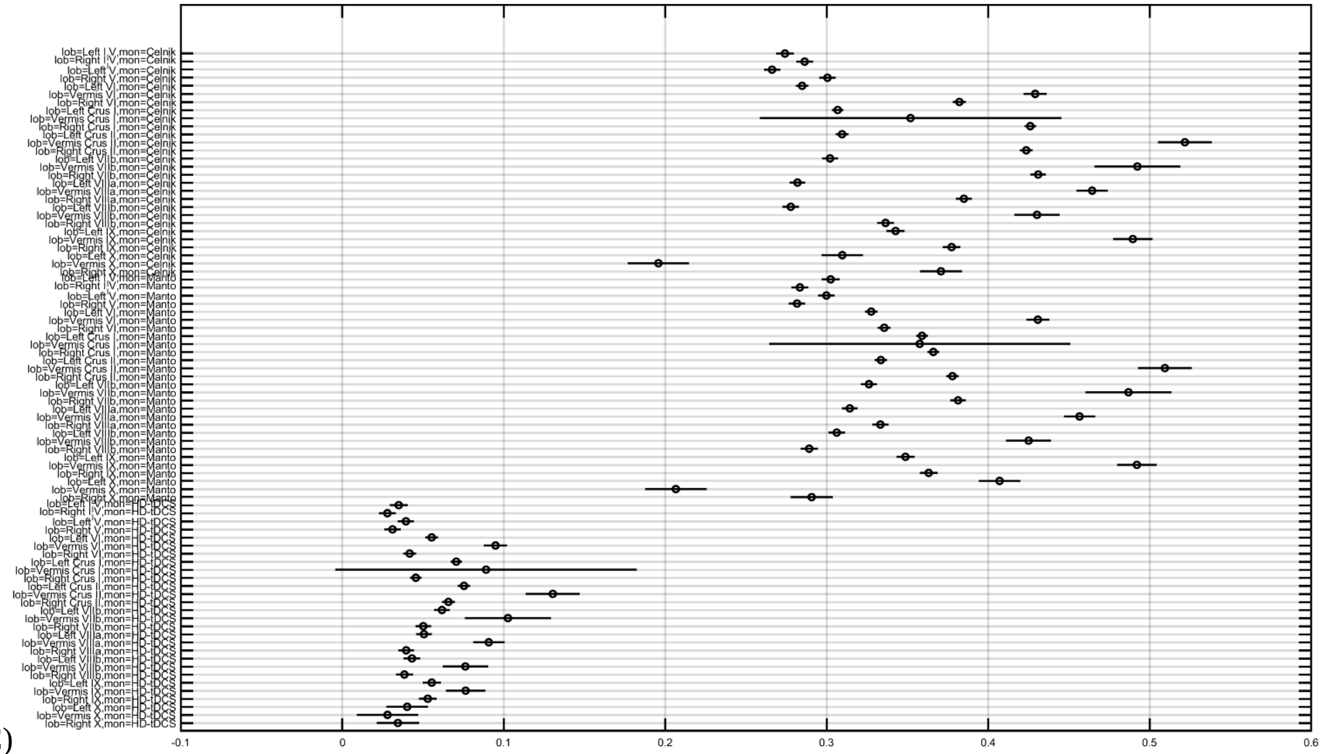


C)

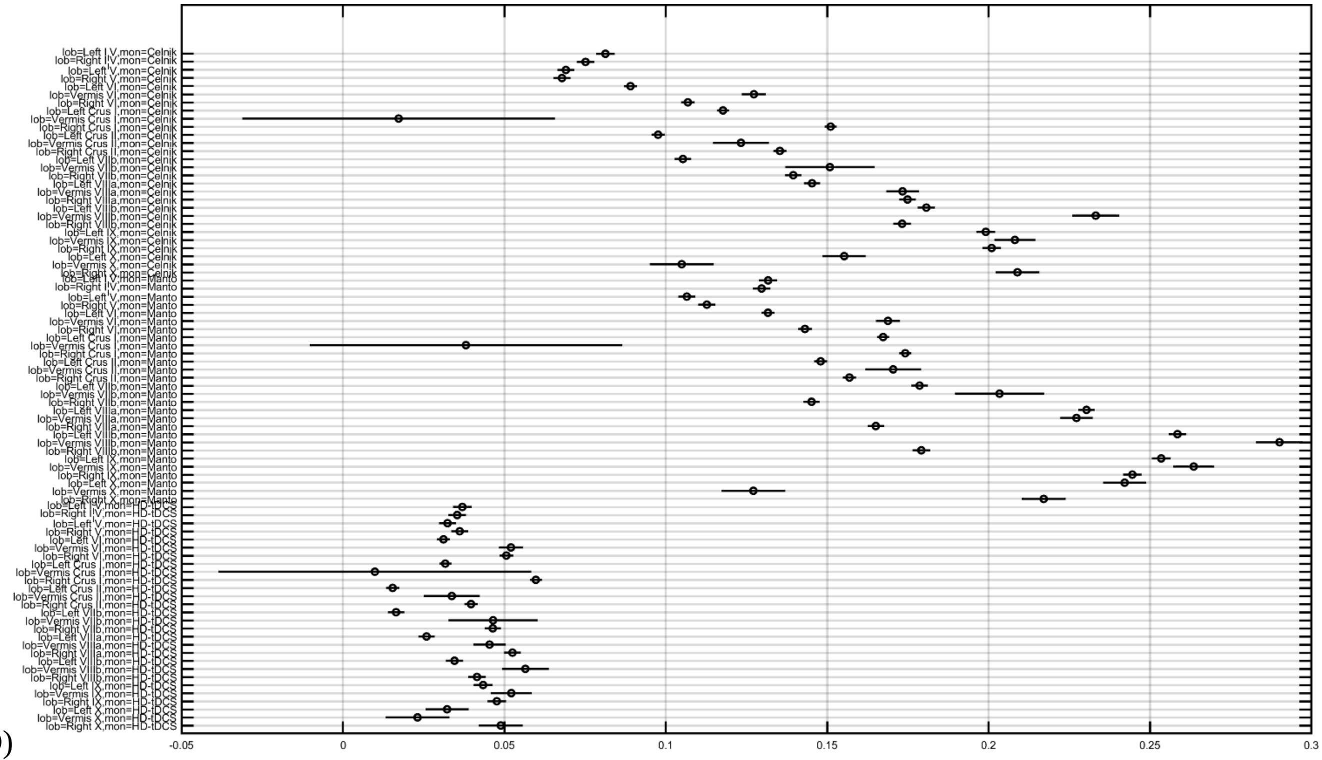


D)

**Figure 1.** Post-hoc multiple comparison with Bonferroni critical values for Celnik, Manto, and HD-ctDCS montages with Colin27 head model for electric field distribution (x-axis is in V/m) with the horizontal lines showing a 95% confidence interval for the true difference of the means (shown with a circle): A) Enorm, B) Ex, C) Ey, D) Ez.

**Supplementary Table 1.** Electric field strength (V/m) mean±standard deviation at cerebellar lobules due to Celnik ctDCS montage

| Cerebellar Location | Left side (V/m) | Right side (V/m) | Vermis (V/m) |
|---|---|---|---|
| | EF mean±std | EF mean±std | |
| I-IV | 0.2930±0.1269 | 0.3206±0.1754 | |
| V | 0.2763±0.0154 | 0.3248±0.2002 | |
| VI | 0.3022±0.1665 | 0.4180±0.2160 | 0.4509±0.0160 |
| Cr I | 0.3351±0.1335 | 0.4698±0.2305 | 0.3762±0.2382 |
| Cr II | 0.3341±0.1162 | 0.4531±0.1795 | 0.5461±0.1153 |
| VIIb | 0.3291±0.0112 | 0.4657±0.1671 | 0.5162±0.0121 |
| VIIIa | 0.3275±0.1356 | 0.4447±0.1975 | 0.5002±0.0721 |
| VIIIb | 0.3405±0.1375 | 0.4003±0.1939 | 0.4961±0.0580 |
| IX | 0.4068±0.1693 | 0.4496±0.1379 | 0.5383±0.0319 |
| X | 0.3590±0.1396 | 0.4982±0.2530 | 0.2308±0.1894 |

**Supplementary Table 2.** Electric field strength (V/m) mean±standard deviation at cerebellar lobules due to Manto ctDCS montage

| Cerebellar Location | Left side (V/m) | Right side (V/m) | Vermis (V/m) |
|---|---|---|---|
| | EF mean±std | EF mean±std | |
| I-IV | 0.3518±0.1533 | 0.3179±0.1690 | |
| V | 0.3347±0.1837 | 0.3047±0.1865 | |
| VI | 0.3740±0.2029 | 0.3669±0.1876 | 0.4694±0.1644 |
| Cr I | 0.4151±0.1661 | 0.4092±0.1995 | 0.3929±0.2484 |
| Cr II | 0.3910±0.1422 | 0.4141±0.1561 | 0.5496±0.1176 |
| VIIb | 0.4012±0.1561 | 0.4130±0.1419 | 0.5346±0.0104 |
| VIIIa | 0.4219±0.1896 | 0.3772±0.1606 | 0.5184±0.0740 |
| VIIIb | 0.4300±0.1866 | 0.3451±0.1637 | 0.5248±0.0638 |
| IX | 0.4563±0.1861 | 0.4484±0.1459 | 0.5689±0.0351 |
| X | 0.5333±0.2084 | 0.3839±0.1889 | 0.2522±0.2044 |

**Supplementary Table 3.** Electric field strength (V/m) mean±standard deviation at cerebellar lobules due to HD-tDCS montage

| Cerebellar Location | Left side (V/m) | Right side (V/m) | Vermis (V/m) |
|---|---|---|---|
| | EF mean±std | EF mean±std | |
| I-IV | 0.0558±0.0254 | 0.0571±0.0328 | |
| V | 0.0539±0.0320 | 0.0632±0.0396 | |
| VI | 0.0646±0.0384 | 0.0903±0.0485 | 0.1157±0.0423 |
| Cr I | 0.0785±0.0328 | 0.1076±0.0555 | 0.0934±0.0595 |
| Cr II | 0.0787±0.0275 | 0.1082±0.0392 | 0.1425±0.0316 |
| VIIb | 0.0657±0.0224 | 0.0929±0.0321 | 0.1199±0.0062 |
| VIIIa | 0.0581±0.0237 | 0.0849±0.0340 | 0.1084±0.0182 |
| VIIIb | 0.0569±0.0233 | 0.0690±0.0345 | 0.1008±0.0119 |
| IX | 0.0735±0.0323 | 0.0819±0.0262 | 0.0967±0.0055 |
| X | 0.0527±0.0208 | 0.0761±0.0375 | 0.0382±0.0309 |
